# Contrasted effects of human pressure on biodiversity in the UK: a multi-taxonomic assessment using airborne environmental DNA

**DOI:** 10.1101/2025.05.01.651649

**Authors:** Orianne Tournayre, Joanne E. Littlefair, Nina R. Garrett, Andrew S. Brown, James J. Allerton, Melania E. Cristescu, Elizabeth L. Clare

## Abstract

Human activities have significantly modified habitats, resulting in a global biodiversity crisis. Here, we leveraged the first national-scale biodiversity survey based on airborne environmental DNA, comparing the effects of three human pressure indices increasing in complexity and scope across diverse vertebrates, insects, plants and fungi. While most taxa exhibited higher diversity in urban areas compared to rural ones, we uncovered more complex patterns using the landscape-pollution and human footprint indices, including dual diversity peaks at both high and moderate levels of human pressure. We also show an effect of human pressure on community composition even when local species richness remained stable: regardless of the human pressure index, anthropogenic sites were mostly characterized by synanthropic and invasive species. Overall, our results underscore the complex interactions among anthropogenic pressures, taxon diversity and community composition, demonstrating the value of multi-taxon analyses and multiple indices to better understand biodiversity patterns at large scales.

## INTRODUCTION

Anthropogenic activities have caused a multifaceted biodiversity crisis on a global scale including loss of species abundance and biomass, altered distribution and extinctions (*1*, *2*). The increased demand on land and resources by the human population has led to significant habitat modification and loss (*3*, *4*). For instance, urban areas have more than doubled since 1992 and global urban cover will likely reach 1.7 × 10^6^ km^2^ by 2050 (*5*). Contrasting effects of urbanization on biodiversity led to competing hypotheses regarding the impact of anthropogenic modifications on taxonomic communities (62 hypotheses, review in (*6*)).

First, cities can be considered as biological deserts, with lower species diversity at their core than non-urban areas. A straightforward explanation is that urbanization fundamentally alters the ecosystem structure and function with the replacement of habitable areas by impervious surfaces (e.g. building), and the degradation of remaining habitat. Beyond the disappearance of critical resources resulting in direct biodiversity loss, there is also an impact on species phenology (e.g. heat island effect on plants; (*7*)), a disruption of species behavior and communication due to traffic noise and light pollution (*8*, *9*) or the creation of barriers to movement (e.g. streets).

In contrast, cities have been hypothesized to be hotbeds of biodiversity with an increasing proportion of non-native and invasive species towards the urban core, contributing positively to the overall species diversity (*10*, *11*). Some native taxa, especially synanthropes and generalists, also contribute to higher richness in cities as their populations benefit from the spatial heterogeneity (e.g. mosaic of impervious surfaces and green spaces), as well as the new nesting and food resources (*12–14*). As urbanization intensifies, urban species assemblages become more similar to each other than the species assemblages of the ecosystems they replace (*15*). This process is well-known as “urban biotic homogenization” and describes species composition similarity across different cities (*16*). Unlike animals, plant diversity and community composition in urban environments are largely shaped by deliberate human actions (e.g. ornamental vegetation, vegetable gardening). However, native species such as early successional plants can also become widespread and contribute to the global homogenization across cities (*17*).

However, species’ response to urbanization can be more complex than a pattern of higher or lower richness in cities. Some studies focusing on a rural-urban gradient have identified peaks of plant and animal richness at intermediate levels of urbanization (i.e. suburbs) (*10*, *18*). These hump-shaped patterns align with a key concept in ecology, stating that species coexistence is the highest when disturbance occurs at intermediate frequency or intensity (“Intermediate disturbance hypothesis”, (*19*, *20*)). This is due to a trade-off between a species’ ability to compete and its ability to tolerate disturbance, thereby favoring the coexistence of competitive dominants and colonizers (*21*).

While these three competing hypotheses have been assessed on smaller scales, a significant gap remains in understanding how anthropogenic pressures shape biodiversity at much larger scales and across a wide range of taxonomic groups. Most studies on human impact have focused on specific taxa, particularly plants and birds, investigating local urban-rural gradients at the landscape and habitat levels (*22*). Biodiversity patterns along these gradients can be complex with variation between cities from the same geographic area (*11*), and within cities (*23*). Moreover, specific land uses (e.g., private and public green spaces) or management practices (e.g. livestock grazing) may play an important role in biodiversity patterns (*24*, *25*). Finally, another challenge lies in the way urban and non-urban areas are contrasted: the proportion of impervious surfaces (*10*, *26*). Although rural areas have lower impervious surfaces, agriculture and farming practices coupled with the extensive use of pesticides have both isolated and reduced the size of suitable habitat patches for wildlife, causing important species loss and homogenization of local communities (*27*, *28*). A comprehensive assessment of the overall effects of land use on biotic communities should therefore encompass urban, agricultural and (semi)natural landscapes at large scale, as well as including a wide variety of taxonomic groups since their presence is interconnected but their responses to anthropogenic disturbances vary (*10*, *29*).

Assessments which are taxonomically broad are rare due to the challenge of conducting large scale biodiversity assessments. However, recent studies have shown that it is possible to survey many phyla simultaneously on a large scale using airborne environmental DNA (eDNA) (*30*, *31*). Similar to other eDNA-based approaches, airborne eDNA approaches bypass the need for capturing or observing individual organisms: in a single sample, airborne eDNA can recover a broad range of taxonomic groups, including elusive, rare, difficult-to-capture species or challenging to identify without highly specialized expertise (*32*). While airborne eDNA offers numerous advantages over field surveys, it also has limitations. Environmental factors affect detection success by degrading and transporting DNA, and sample processing (e.g. DNA extraction, PCR) also introduces biases preventing individual organism counts. However, a key advantage of airborne eDNA is its ability to leverage existing infrastructure, such as national air quality monitoring networks (*30*, *31*). Continuous sampling from such infrastructure already deployed for other environmental monitoring purposes, enables national-scale biodiversity assessments, offering broad spatial and temporal coverage without the logistical challenges and costs typically associated with traditional field surveys (*31*).

In this study, we assessed the impact of human pressure on terrestrial biodiversity at a national scale, and across various taxonomic groups including mammals, birds, insects, plants and fungi. We investigated both alpha and beta diversity patterns across the country using three human pressure indices increasing in complexity and scope. The first index, developed by the Heavy Metals air quality network in a human health context, classified the sites as either urban or rural based on their proximity to built-up areas. The second index combined natural and anthropogenic land cover data with the concentration of particulate bound heavy metals in the air (pollution emitted by traffic and industrial processes as a proxy of human activity) to reflect a gradient of human pressure beyond the binary urban-rural distinction. The last index, the global human footprint index, combines eight sub-indices representing different aspects of human pressures, including population density, infrastructures and anthropogenic land covers (*33*). We hypothesised that 1) biodiversity patterns would be influenced by all three human pressure indices, and 2) different taxonomic groups would show contrasted responses to human pressure. Specifically, we predicted that alpha diversity would decline from rural to urban sites across all taxonomic groups, with the exception of plants, from which we expected to increase due to the introduction of diverse non-native species for ornamental purposes in public and private green spaces. In contrast, when considering the two gradient-based indices, we anticipated that alpha diversity might follow a hump-shaped response, peaking at intermediate disturbance levels. Lastly, we predicted that beta diversity would decrease under increasing human pressure, reflecting biotic homogenization across all taxonomic groups.

## RESULTS

### 1. Land cover - pollution index: determining PCA axes explaining the most variation

The first PCA axis (PC1, 41.8%) represented a gradient running from high to low levels of heavy metal pollution, and the second PCA axis (PC2, 16.6%) separated the sites characterized by anthropogenic landscape (agricultural areas, artificial surfaces) to those characterized by natural and semi-natural elements (Figure 1). The urban sites were situated in the anthropogenic land cover and polluted section (except Belfast Centre and Swansea Coedgwilym), and the rural sites were clustered by the absence of pollutants in more or less anthropogenic landscapes (Figure 1).

**Figure 1.**
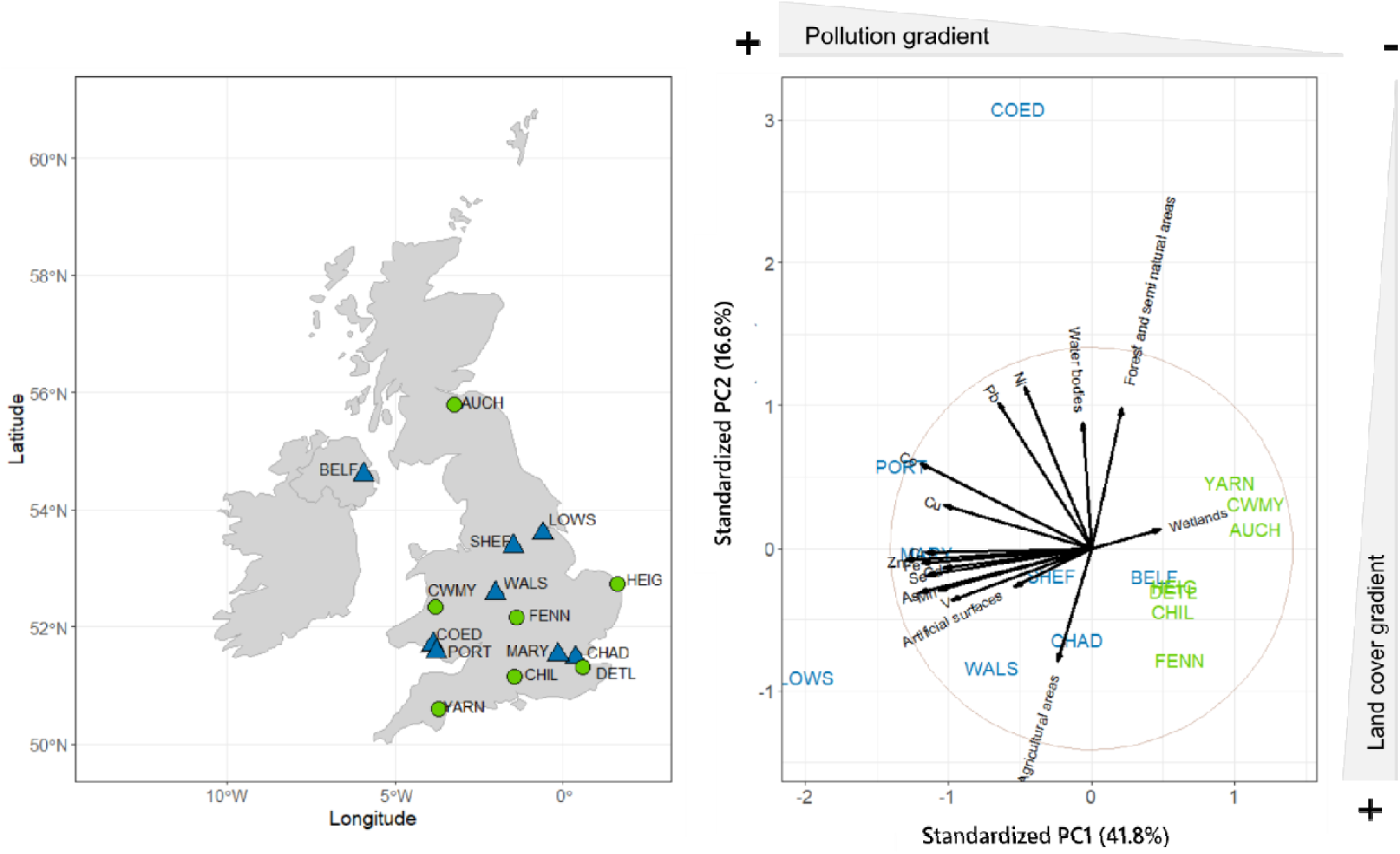
Characterization of the sites. Using A) the environment types as defined for the Heavy Metals Network (Urban = blue triangles, Rural = green circles) (left panel), and B) a Principal Component Analysis combining five CORINE Land Cover features within a radius of 18.6 km around the sites (waterbodies, forest and semi-natural areas, wetlands, artificial surfaces and agricultural areas) and the heavy metal concentrations recorded by the heavy metals air quality network in 2022 (right panel): arsenic (As), cadmium (Cd), cobalt (Co), copper (Cu), iron (Fe), manganese (Mn), nickel (Ni), lead (Pb), selenium (Se), vanadium (V) and zinc (Zn). Rural sites were represented in green and urban in blue, on both panels. BELF = Belfast Centre, AUCH = Auchencorth Moss, LOWS = Scunthorpe Low Santon, SHEF = Sheffield Devonshire Green, WALS = Walsall Pleck Park, HEIG = Heigham Holmes, FENN = Fenny Compton, MARY = London Marylebone Road, CHAD = Chadwell St Mary, DETL = Detling, CHIL = Chilbolton Observatory, YARN = Yarner Wood, CWMY = Cwmystwyth, COED = Swansea Coedgwilym, and PORT = Port Talbot Margam. The latitude and longitude of each site are publicly available on the UK AIR Air Information resource website (https://uk-air.defra.gov.uk/interactive-map?network=metals).

### 2. Alpha diversity

Our refined dataset, with samples collected in 2022 and focusing on specific taxonomic groups (birds, mammals, insects, plants, fungi), comprised a total of 12,443,740 reads and 1,213 ASVs assigned to unique taxa. Similar to the original dataset (*31*), we identified endangered species (e.g. European hedgehog), ornamental and cultivated plants (e.g. squash, geranium, fuchsia, cypress), invasive (e.g. Reeve’s muntjacs and Eastern grey squirrel), commensal (mice, rats) and charismatic species (e.g. badgers, red fox, bats), as well as vectors of diseases (e.g. mosquitoes). We reached or almost reached the accumulation curve plateau for most taxonomic groups at the site level (Figure S1).

#### 2.1. Effect of ambient storage time and read counts

Ambient storage time of the filters used to generate the biodiversity data had a negative effect on the richness of all taxonomic groups, regardless of the human pressure index (Tables S1 to S3). In contrast, read count had a positive effect on all taxonomic groups but insects using the Urban-Rural index (Table S1) and the Human Footprint index (Table S3), and a positive effect on bird, mammal and plant richness using the Land cover - Pollution index (Table S2).

#### 2.2. Urban vs Rural index

Richness diversity at the sites reached 32 bird species (Scunthorpe Low Santon), 14 mammal species (Walsall Pleck Park), 98 insect genera (Scunthorpe Low Santon), 34 plant genera (Walsall Pleck Park) and 35 fungi genera (Walsall Pleck Park). All taxonomic groups showed lower richness in rural than urban environments (Figure 2A and Table S1), except fungi (E = 0.223, p = 0.180) showing no significant difference.

**Figure 2.**
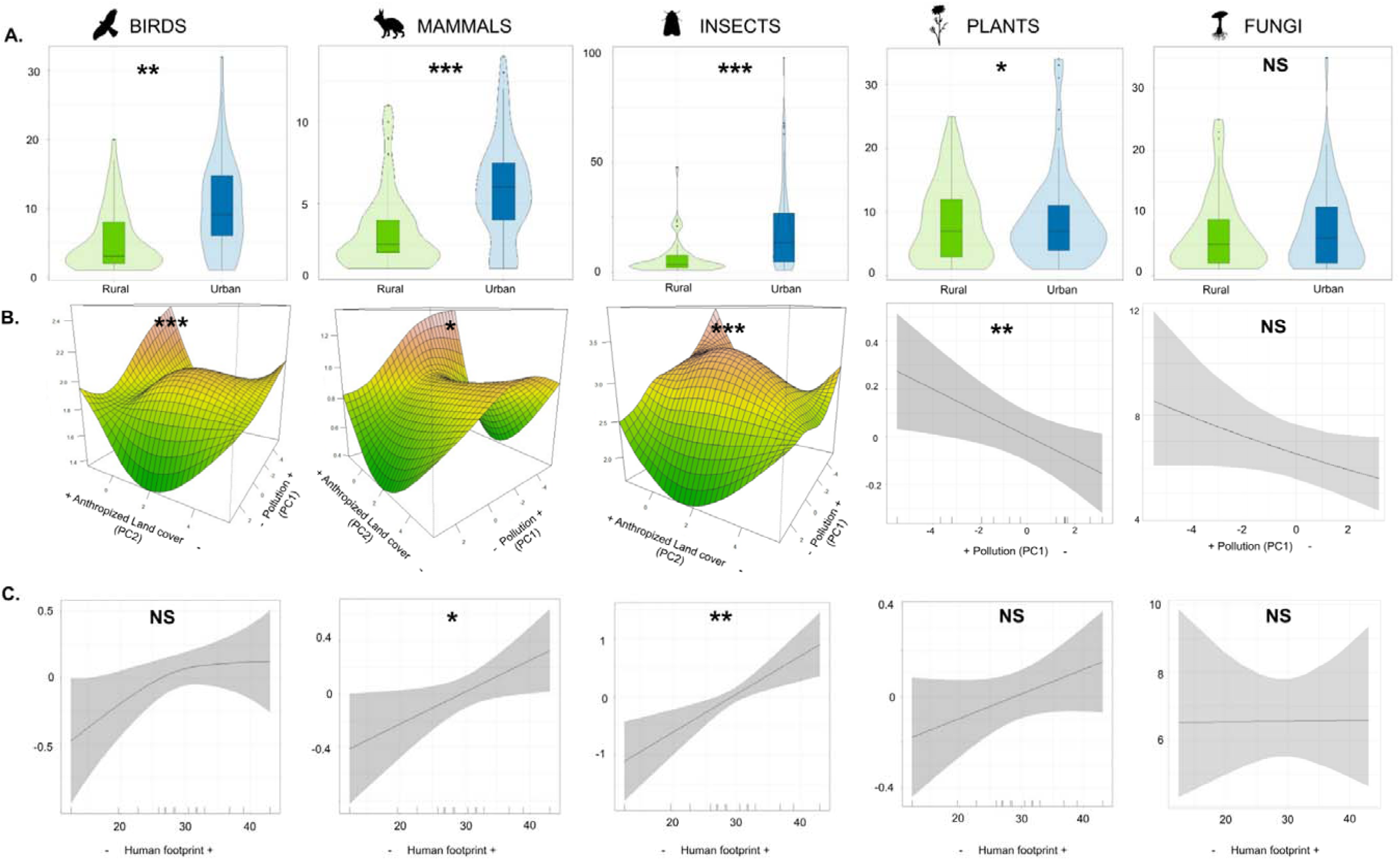
Effect of human pressure on taxon richness for each group of taxa. Human pressure was defined as **A)** the binary environment e index based on distance to built-up, with rural in green (N = 7 sites) and urban in blue (N = 8 sites). The y-axis indicates the richness ues. The violin plots behind the boxplots show the density of the data.; **B)** the land cover-pollution index composed of a gradient of heavy tal pollution (PC1, from the highest concentrations indicated by a “+” to the lowest concentrations indicated by a “-”) and land cover (PC2, m the most anthropogenic indicated by a “+” to the least anthropogenic indicated by a “-”). For birds, mammals and insects, the surface apes were obtained with the mgcv::vis.gam function and represent the smooth interaction term of the GAMM model, indicating how the ationship between PC1 and the predicted richness depends on the value of PC2, with the y-axis indicates the estimated value from the model ear predictor scale). For plants, the GAMM plots estimated smooths of the significant land cover-pollution variables were obtained with the tia::draw function with the Y-axis representing partial effects. For fungi the best model was a GLMM and as only the effect of PC1 was nificant, only PC1 is represented here (ggeffects::ggpredict function; y-axis is on the response scale); **C)** the human footprint index. For all onomic groups but fungi, the GAMM plots estimated smooths were obtained with the gratia::draw function with the Y axis representing partial ects. For fungi the best model was a GLMM (ggeffects::ggpredict function; y-axis is on the response scale). The significance of the p-values each facet is indicated as follow: * for p < 0.05; ** for p < 0.01, *** for p < 0.001 and not significant (NS) for p > 0.05. “Mammals” does not lude domestic mammal species.

#### 2.3. Land cover - pollution index (LPI)

Plant richness showed a decrease along PC1 (i.e. increased richness with air pollutant concentration; Figure 2B), while fungi richness was not influenced by either component of the LPI (Table S2). In contrast, a significant non-linear interaction between PC1 and PC2 was observed for birds, mammals (excluding domestic species) and insects. For all three taxonomic groups, richness was highest in the most anthropogenic environments - corresponding to low values of both PC1 and PC2 (Figure 2B; Table S2) - and then followed a saddle-shaped pattern: in areas with highly anthropogenic land cover, richness increased along the pollution gradient, whereas in low to intermediately anthropogenic land cover, richness peaked at intermediate levels of pollution (Figure 2B; Table S2). In contrast, when domestic species were included, mammal richness was influenced only by the pollution gradient PC1, plateauing in polluted sites (Table S2).

#### 2.4. Human footprint index (HFI)

We did not detect any significant impact of the human footprint on bird, plant and fungi richness (Figure 2C; Table S3). However, we observed a significant increase in mammal and insect richness along the human footprint gradient (Figure 2C; Table S3).

### 3. Beta diversity

The indicator species analysis identified species whose presence was significantly associated with either rural or urban environments. Rural environment was associated with a compact tufted perennial grass (*Cynosurus* genus), while urban environment was associated with four common bird species (e.g. European starling, Eurasian collared dove, Eurasian sparrowhawk, grey heron), three domestic/commensal/invasive mammal species (e.g. domestic cat, wood mouse, Reeves’s muntjac), 14 various insect genera, and two fungi genera (Table S4).

Total beta diversity values partitioned by environment type (rural and urban) were similar (Δ_rural-urban_ ≤ 0.08), with slightly higher total beta diversity in rural environments across all taxonomic groups, except for plants (Figure S2). Beta diversity was primarily determined by turnover in all taxonomic groups and environment types (Figure S2). We observed an effect of the environment type on bird (R² = 0.121, p = 0.005), mammal (R² = 0.160, p = 0.002; R²_with domestic species_ = 0.179, p_with domestic species_ = 0.003) and insect (R² = 0.105, p = 0.002) communities (Figure 3, Table S5A). When considering the land cover - pollution index, we observed 1) different communities of birds (R² = 0.115, p = 0.029), mammals (only when including domestic species; R² = 0.119, p = 0.037) and insects (R² = 0.098, p = 0.002) along the pollution gradient (PC1 axis), and 2) different fungi communities along the land cover gradient (PC2; R² = 0.094, p = 0.031) communities (Figure 3; Table S5B). Finally, when quantifying human pressure with the human footprint index, we observed an effect on bird (R² = 0.150, p = 0.001), mammal (R² = 0.183, p = 0.001; R²_with domestic species_ = 0.195, p_with domestic species_ = 0.001), insect (R² = 0.103, p = 0.003) and plant (R² = 0.091, p = 0.019) community dissimilarities (Figure 3; Table S5C).

**Figure 3.**
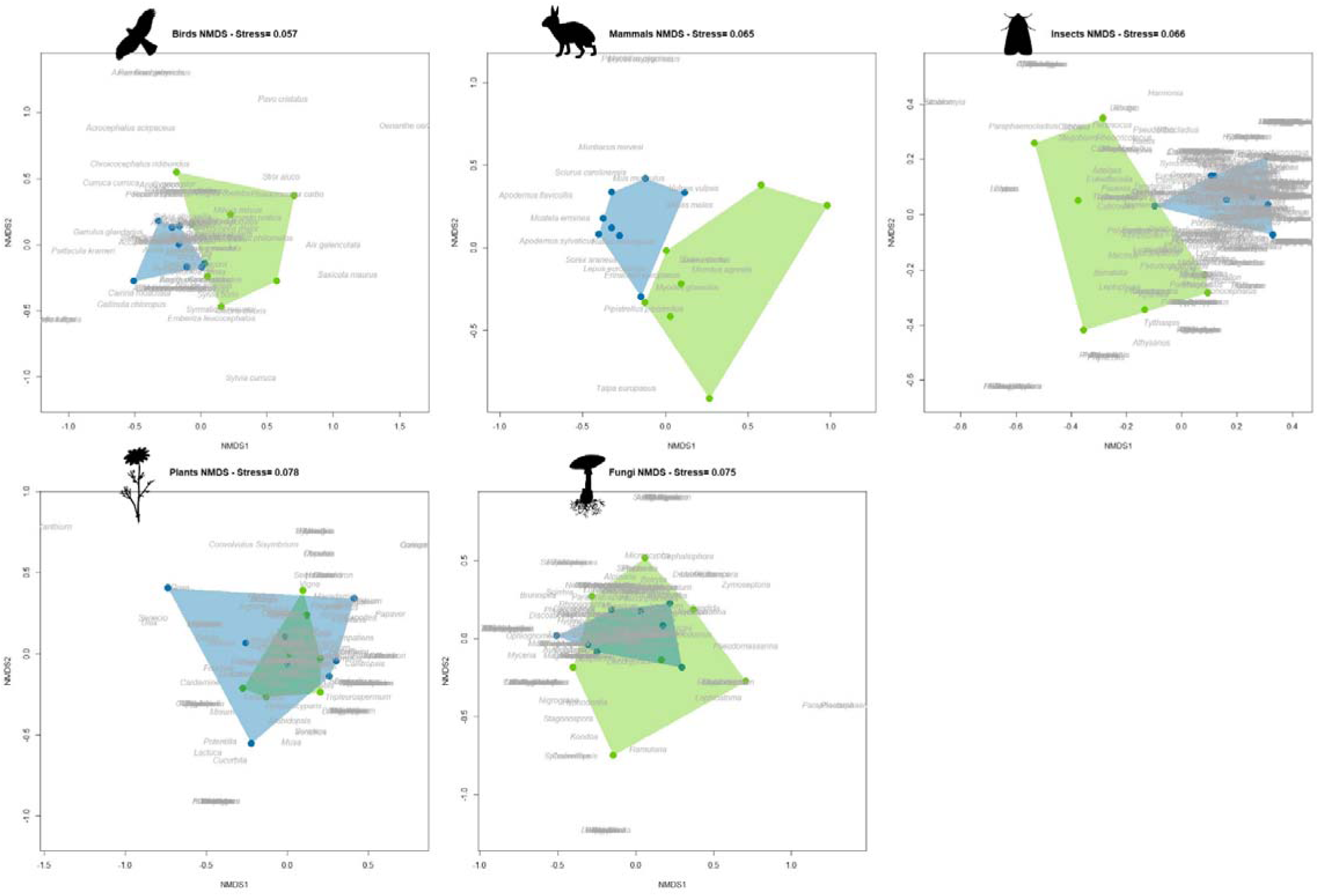
NMDS representation of community dissimilarity (Jaccard). Dot represents the sites (green = rural, blue = urban). Species (birds, mmals without domestic species) and genus (insect, plant, fungi) names are indicated in grey.

## DISCUSSION

We found diverse alpha diversity patterns across taxonomic groups and human pressure indices, highlighting the importance of considering diverse indices and non-monotonic relationships when assessing human impacts on biodiversity (*34*, *35*). A simple binary urban-rural classification was too coarse. For example, we detected lower bird, mammal, insect and plant richness in rural environments compared to urban ones, and no significant differences were found for fungi. However, when considering a broader spectrum of land covers and pollutants as a proxy of human activity (land cover-pollution index; LPI), the relationship between richness and human pressure was more complex and taxonomic-specific. While fungal richness was not influenced by either component of the land cover-pollution index, plant richness increased with human activity (pollutant concentrations). This pattern aligns with previous studies suggesting that cities are hot spots of plant species richness (review in (*36*)), likely because of a combination of factors such as a high prevalence of ornamental species, niche expansion and increased carrying capacity of cities associated with their habitat heterogeneity (*10*, *36*, *37*). This may in turn enhance insect richness by providing favourable habitat and resources (*38*). For example, ornamental plants are known to attract a broad spectrum of flower-visiting insects (*39*), and (*40*) showed that higher bee species richness is supported in urban areas than agricultural land in the UK. In our study, insect richness overall mirrored that of plants but two distinct relationships were observed depending on the human pressure index used. Using the human footprint index, insect richness increased with human pressure. Using the land cover-pollution index, insect richness - like birds and mammals (excluding domestic species) - responded to both human activity (pollution gradient PC1) and land cover (PC2) through a non-linear, saddle-shaped interaction (Figure 2). Specifically, richness was highest in the heavily anthropogenic and polluted sites, with a secondary peak at intermediate levels of human pressure. Our dataset included numerous species, beyond domestic animals, known to live in close proximity to humans for food and/or roosting resources (e.g. pigeons, rodents, red fox, pipistrelle bats) that could have driven this pattern. This suggests that while heavily human-modified areas may offer novel niches and abundant resources, moderate levels of human pressure may also support high biodiversity, possibly due to a balance between disturbance and habitat availability.

This dual pattern provides a nuanced perspective that integrates two commonly proposed hypotheses in urban ecology: cities as biodiversity hotspots (*10*, *11*) and the intermediate disturbance hypothesis (e.g. (*10*, *34*, *41*)). Rather than being mutually exclusive, our findings suggest that these hypotheses could co-occur within and across broad taxonomic groups, likely reflecting diverse species-specific patterns (e.g. urban avoiders, urban exploiters, (*42*)). Moreover, insect, plant and fungi were identified at the genus-level only, potentially masking some divergent trends that would lead to generalized patterns that may not accurately reflect group-specific responses (*34*, *43*). Future study with enough statistical power per taxonomic unit should investigate intra-group patterns for a refined understanding of human pressure on biodiversity. Interestingly, when including domestic mammal species to the analysis, we found that the interaction between the two components of the land cover - pollution index was no longer significant and mammal richness responded only to the pollution gradient— showing a plateau of high diversity at high pollution levels (i.e., areas of intense human activity). While domestic animals are associated with human presence and management, they participate in ecological networks and can influence local biodiversity patterns. For example, cats intensively impact wildlife mostly through predation and competition (review in (*44*)), while livestock (e.g. horse, goats, cattle) can have both positive and negative effects on plant and invertebrate communities (review in (*45*)). This raises important questions about how we define and measure biodiversity in anthropogenic environments. Including domestic species in analyses may obscure patterns specific to wild species, but excluding them entirely could overlook important drivers of species interactions and community composition in anthropogenic landscapes.

While local diversity, i.e. alpha diversity, has been the focus of many studies, human pressures also induce significant changes in species assemblages (*34*). Beta-diversity provides a measure of community dissimilarity and can be partitioned into two additive components: the turnover accounting for the dissimilarity due to the replacement of some species by others between assemblages, and the nestedness accounting for the dissimilarity associated with species losses in which an assemblage is a subset of another (*46*, *47*). While we found no effect of geographical distance among our 15 sites across the UK, we identified a strong dominance of turnover in beta diversity for all taxonomic groups, indicating that species replacement was a key factor in shaping community composition in the country (*48*, *49*). However, while total beta diversity was similar among urban and rural sites, the community composition of the urban sites was slightly more homogenised (i.e. lower beta diversity). Biotic homogenization is a well-known process in studies on human impact on biodiversity (*16*), with the replacement of specialist species by generalist ones (*17*, *50*). This aligns with our findings, as in our study anthropogenic environments were characterized by urban adapters and exploiters (*18*, *42*), i.e. common, cosmopolitan and generalist taxa such as starlings, mice, and invasive species (e.g. muntjac). Interestingly, the indicator species analysis showed many taxa associated with urban environment and only one with rural environment. This pattern underscores how biotic homogenisation results in urban communities composed of a consistent subset of generalist and human-associated taxa, while rural areas are more heterogeneous in composition. Overall, species assemblages were impacted by human pressures, with animal community dissimilarity more consistently impacted across the three human pressure indices, while plant community dissimilarity was only influenced by the human footprint index, and fungi dissimilarity only by the land cover gradient. These findings highlight the importance of examining both alpha and beta diversity when assessing the ecological impact of human activity. As changes in community composition can occur despite stable local richness, focusing solely on species richness could overlook significant compositional changes that have critical implications for conservation and ecosystem functioning (*51*, *52*). Although the scope of our study focuses on human impact on biodiversity, the relatively moderate variance explained by the models (min_R²_ = 4.2%, max_R²_ = 19.5%) suggested that other factors, such as climate, soil properties, or canopy characteristics (*53*, *54*), might contribute to the observed community dissimilarities.

Collecting diversity data across a national scale using traditional survey approaches would have been challenging for such a broad range of taxa as it requires diverse taxonomic expertise, as well as significant financial and logistical resources. Biomonitoring using airborne eDNA from air quality monitoring networks has recently been brought to the forefront as a very promising and scalable approach (*30*, *31*). As the infrastructures filter the air almost continuously, the recovered diversity patterns encompass species phenology and sampling timing. In this study, we were relatively limited in our number of sites (N = 15), with a greater concentration of sites in southern England and Wales, and only a single site in Northern Ireland and Scotland. These constraints arose from the opportunistic use of filters initially collected for heavy metal analysis. Nonetheless, the sampled sites encompass a diverse range of habitats, providing an important opportunity for broad biodiversity assessment and human pressure evaluation. While taxa recovery was negatively influenced by filter storage conditions (initially collected for heavy metal analyses and therefore not stored for DNA analyses) and positively influenced by read count, we were able to detect the effect of human pressure on diversity. However, further studies with eDNA-optimized storage protocols and a more balanced design across the country will enable a finer resolution and generalization of the observed biodiversity patterns at the national scale.

The ecology of airborne eDNA, including its dispersal and deposition, is still not well-known, but is likely to be influenced by physical and biological factors that impact our ability to detect eDNA signals. Rapid particle deposition from the air limits the time window for detecting a species, which provides the great advantage of a real-time snapshot of species presence but may also increase the rate of false negatives (i.e., species present but not detected if deposition happened too quickly). For example, similar to particulate matter, rain could wash eDNA out of the air (*55*). Moreover, while wind can be considered as a conveyor of biodiversity by transporting eDNA, it also complicates the exact attribution of source and the spatial resolution at which human pressure can be assessed. For example, urban areas with dense buildings could create complex wind patterns potentially trapping eDNA locally and leading to an overestimation of diversity metrics (e.g. inflated richness). On the other hand, their higher levels of particulate matter in the air (e.g. from vehicle emissions) could act as PCR inhibitors, thereby reducing our ability to recover eDNA. In this study, we used a buffer of 18.6 km around each site based on the estimated median eDNA transportation distance (*31*), to calculate the land cover-pollution and human footprint indices. While it is in the range of previous suggestions (≤100 km; (*56*, *57*)), it is possible that some eDNA sources are located further away, as smaller particles included in the PM_10_ fraction sampled here can travel long distances (e.g. some PM_2.5_ pollution in the UK originates from continental Europe; (*58*)). Thus, the spatial scale may not accurately reflect the most relevant habitat range for all taxonomic groups. For instance, while such a broad buffer may be appropriate for wide-ranging taxa like birds or mammals, it could overlook finer-scale habitat use by less mobile organisms such as many insects or plants. Future research must therefore assess the influence of particle size, shape (e.g., aerodynamic properties), and environmental factors (e.g., wind speed, wind direction, rainfall, topography) on detection success and spatio-temporal resolution to refine the observed patterns. Finally, as with other eDNA-based metabarcoding studies, DNA extraction efficiency or PCR biases prevents a correlation between sequence and individual abundances (*59*). The relationship between diversity and abundance can be complex as human pressure can impact both or only abundance (e.g. short-term pressure) (*60*, *61*). Thus, further methodological advancement, such as the development of models (*62*), will be invaluable in the next years to move towards quantitative eDNA datasets and estimation of species abundance for biodiversity conservation (*63*).

In conclusion, our study showed strong evidence of human impact on terrestrial biodiversity in the UK, affecting both diversity and community composition. Our multidimensional approach emphasizes the need to consider several human pressure indices and non-monotonic patterns to better capture taxon-specific responses. While our results revealed higher diversity in anthropogenic areas, this does not necessarily imply that populations in such environments are healthy and self-sustaining as urban areas can act as ecological traps (i.e. attractive habitat reducing fitness) (*64*, *65*).

## MATERIAL AND METHODS

### 1. Taxonomic dataset

We used a dataset generated from 185 airborne environmental DNA samples collected between September 2021 and October 2022 at 15 sites of the national United Kingdom heavy metals air quality monitoring network (see (*31*) for detailed information on data generation). Briefly, the instruments sample PM10 onto a cellulose ester filter matter on a weekly basis (*66*). The filters, initially collected for heavy metal analyses, have been stored in Petri dishes (separated by location and month of sampling, resulting in monthly samples) and kept in light-proof cupboards in ambient conditions up to 22 months before being subsampled for DNA analysis and stored at −80°C. The 185 filters along with multiple controls from the field to the sequencing, were processed using a metabarcoding approach with five primer pairs on four genes, including technical replicates to ensure data quality. The resulting dataset comprised 18,373,005 reads belonging to 1,556 ASVs identified to unique taxa across vertebrates (birds, mammals, fish, amphibians), invertebrates (e.g. annelids, arthropods, nematodes), plants (e.g. trees, flowering plants, grasses, algae), protists, and fungi (e.g. microscopic or fruiting-bodies fungi), all spanning a broad range of habitats (terrestrial, sub-terrestrial, semi-aquatic, aquatic). Here, we refined this dataset by keeping only samples from 2022 since data for 2021 were sparse (only a few months), therefore ambient storage time (duration in months the sample was stored at ambient temperature prior to storage at −80°C, (*31*)) ranged from nine to 18 months. We focused on the following terrestrial and semi-aquatic (i.e. at least one aquatic life stage or exploiting aquatic habitats for feeding, or reproduction) taxonomic groups: birds (except exotic caged-birds: finches, budgerigars, cockatiel and grey parrot), mammals (including rabbit, sheep, horse, donkey, cat, dog, goat and pig), insects, non-algae plants, and fungi (except two phyla represented by three ASVs unfrequently detected in the dataset). We visually verified that the sampling effort were sufficient to capture the diversity of each taxonomic group by generating accumulation curves with the specaccum function (1,000 permutations, method= “random”) of the vegan package version 2.6-6.1 (*67*).

### 2. Human pressure indices

#### 2.1. Urban vs rural index

We used the environment type of the UK heavy metals air quality monitoring sites (https://uk-air.defra.gov.uk/networks/site-types). Briefly, a rural station is located at a site “in small settlements and/or areas with natural ecosystems, forests or crops, always more than 20 km away from agglomerations and more than 5 km away from other built-up areas, industrial installations or motorways or major roads”. An urban station is situated at a site in a “continuously built-up urban area”, a traffic station is situated at a site “nearby traffic (roads, motorways, highways), at least 25 m from the edge of major junctions and no more than 10 m from the kerbside”, and an industrial station is situated at a site in the nearest residential area from the industrial area (e.g. power generation, incinerators and waste treatment plants). Five sites belonged to the “Urban” category, one site to “Traffic station”, two sites to “Industrial station” and seven sites as “Rural”. For this study, we considered urban, traffic and industrial stations as “Urban” (Figure 1).

#### 2.2. Land cover - pollution index (LPI)

First, we extracted the summed area surface for each Corine land cover (*68*) in QGIS Desktop 3.38.1 using a buffer zone of 18.6 km around each site (median eDNA transportation distance as estimated in (*31*)), with the limitations of this estimate discussed in the Discussion section. Then we merged them into five main categories: “Artificial surfaces” (111, 112, 121 to 124, 131 to 133, 141 and 142), “Agricultural areas” (211, 222, 231, 242 and 243), “Forest and seminatural areas” (311 to 313, 321 to 324, 331 and 333); “Wetlands” (411, 412, 421 and 423); and finally “Water bodies” (511, 512, 522 and 523). Second, we downloaded the pollutant data (concentrations in ng/m³) available for the year 2022 using the DEFRA UK AIR Air Information Resource website (https://uk-air.defra.gov.uk/data/) (*69*). The data included metals primarily emitted from the combustion of fossil fuels and industrial processes: arsenic, cadmium, cobalt, cooper, iron, manganese, nickel, lead, selenium, vanadium and zinc. Concentrations inferior to the limit of detection were assigned to the upper bound of their range to distinguish them from real “0” values (e.g. <0.9 was set to 0.9). We performed a Principal Component Analysis (PCA) on the land cover and averaged by site heavy metals data using the prcomp function (scale = T, center = T) of the stats v3.6.2 package (*70*). The PCA coordinates on the two axes explaining the most variance were extracted for each site and further used as the land cover - pollution index components.

#### 2.3. Human footprint index (HFI)

The human footprint index dataset was obtained from (*33*) and comprises eight aspects of human pressures to the terrestrial surface: extent of the built environment, population density, electric infrastructure, agricultural lands, pasture lands, roadways, railways, and navigable waterways. The index ranges from 0 to 50, from low to high human pressure. We extracted the average value of aggregated raster cells of the 2013 human footprint maps from (*33*) with a buffer zone of 18.6 km (see above). The HFI overlaps with the LPI described above, as both indices include land cover, but they differ in two main ways: i) the HFI only includes human infrastructures and human-modified land covers, while the LFI includes both natural and semi-natural habitats, ii) the LPI includes heavy metal concentrations in the air as a proxy of human activity, while the HFI includes population density.

### 3. Diversity analyses

#### 3.1. Alpha diversity

We computed taxa richness at the species level for birds and mammals (with and without the domestic species: rabbit, sheep, horse, donkey, cat, dog, goat and pig), and at the genus level for insects, plants and fungi. The analyses were not performed on the Shannon and Simpson alpha diversity metrics, as these metrics rely on the assumption that metabarcoding read counts reflect relative abundance of the taxa, which may not hold with such broad taxonomic coverage due to biases in DNA extraction, PCR amplification or sequencing (*59*).

For each taxonomic group, we performed a generalized linear mixed model fitted with a negative binomial family (GLMM; glmmTMB R package, (*71*)) to evaluate the effect of ambient storage time in months (duration that the sample filter was stored at ambient temperature from collection in 2022 for heavy metal analysis until transfer at −80°C storage for DNA analysis), sequencing depth (read count scaled with the scale function) and human pressure defined as environment type (rural vs urban) on alpha diversity. We included a random intercept for site and for season (Winter: December, January, February; Spring: March, April, May; Summer: June, July, August; Fall: September, October, November) to control for variability in richness due to site-specific environmental conditions and season. The temporal variable “Month of sampling” was not included in the model as it was confounded with “ambient storage time” (one unique value of ambient storage time per month). The absence of temporal autocorrelation was further visually investigated using the acf function. For human pressure defined as land cover-pollution gradient, i.e. the site coordinates on the two first axes of the land cover-pollution PCA (PC1, PC2 and their interaction), we also tested a generalized additive mixed model (GAMM; mgcv R package, (*72*)) fitted with a negative binomial family. We compared GLMM and GAMM models because they assume different relationships between the predictors and response variables: the selection of the best model was done according to the lowest AIC or highest R² in case of ΔAIC < 2 (Table S6). If the interaction between PC1 and PC2 (land cover-pollution variables) in the best model was not significant, we refitted a simpler version without this interaction (Table S6). The effects of the variables were visualized using the vis.gam, draw and the ggpredict functions of the mgcv, gratia (*73*), and ggeffects (*74*) R packages, respectively. The same approach was followed with human pressure defined as the human footprint index.

### 3.3. Beta diversity

We merged the monthly samples by sites to limit the negative effect of ambient storage time on taxa recovery and the effect of a sparse matrix (many rare species). We explored the correlation between taxa presence and the environment type (urban vs rural) for each taxonomic group using the multipatt function of the R package “indicspecies” (*75*) with 999 permutations and func = “r.g”. Using the *beta.div.comp* function (coeff =”BJ”, Baselga Jaccard-based) of the package adespatial (*76*), we identified the relative contributions of the two beta diversity components: dissimilarity due to turnover (i.e. the loss of species is counterbalanced by the gain of new others) and due to nestedness (i.e. the extent to which the composition of one site is a subset of the composition of another site) (*46*). We verified that community composition between sites was not correlated with geographic distance (geodesic) using Mantel tests with the mantel function from vegan (999 permutations; Table S7). Dissimilarities between communities were visualized using NMDS plots with the metaMDS function from vegan (method = “jaccard”, k = 4). The quality of the solution was evaluated based on the stress value (*77*). The effect of the three human pressure indices on community dissimilarities was investigated with a permutational multivariate analysis of variance (perMANOVA; 999 permutations) using the adonis2 function.

## ACKNOWLEDGMENTS

None.

## Author contributions

O.T. designed the study with contributions from E.L.C, J.E.L and M.E.C. O.T. performed the analyses and wrote the draft of the manuscript, and all authors contributed to edits.

## Data and materials availability

The datasets and R code used to generate the results will be publicly available on the FigShare repository upon acceptance of the manuscript.

## SUPPLEMENTARY FIGURES

**Figure S1.**
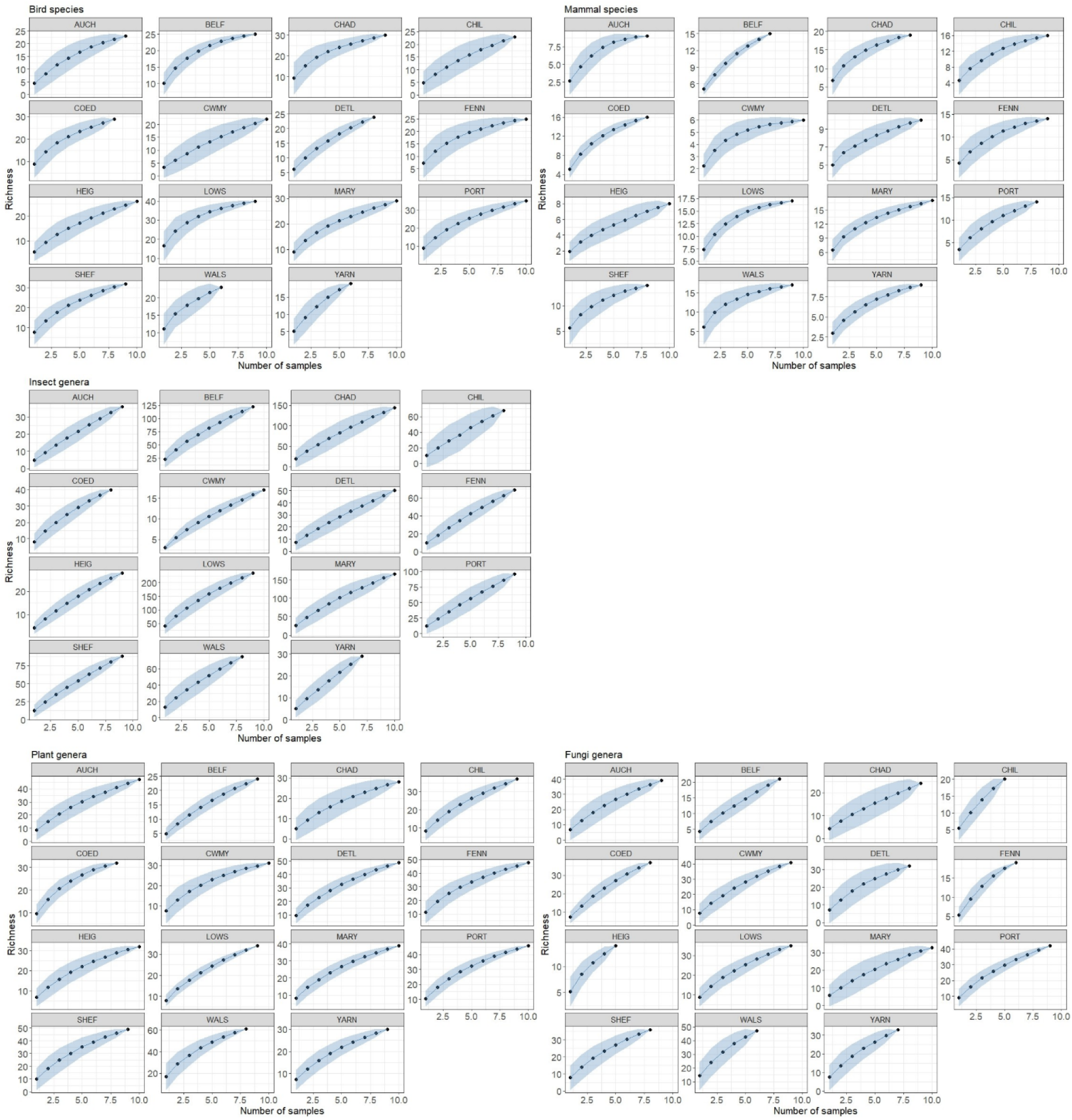
Richness accumulation curves for each site and taxonomic groups. Taxonomic groups include bird species, mammal species, insect genera, plant genera and fungi genera. The dots represent the richness value and the shaded area the standard deviation, as obtained with the specaccum function of the vegan package (1,000 permutations).

**Figure S2.**
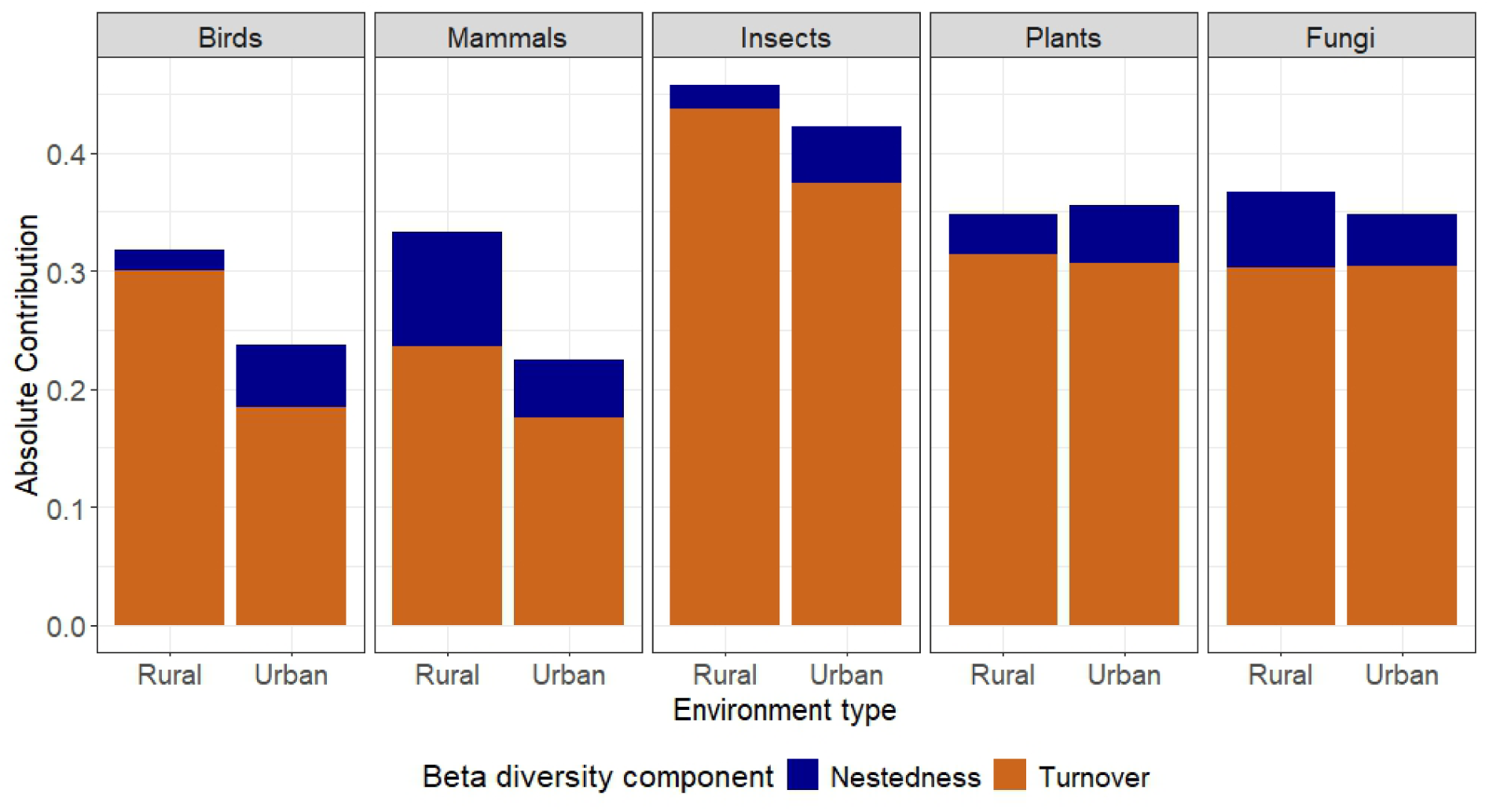
Contribution of turnover and nestedness to total beta diversity. Contribution of turnover and nestedness to total beta diversity (BDTotal) using the Jaccard-based index of Baselga family in rural and urban sites for each taxonomic group (bird species, mammal species, insect genera, plant genera and fungi genera). Rural = 7 sites, Urban = 8 sites. “Mammals” does not include domestic mammal species.

## SUPPLEMENTARY TABLES

**Table S1.** Outputs of the GLMM investigating the effect of ambient storage time, read counts and human pressure defined as the urban - rural index on alpha diversity (richness). E = Estimate, SE = Standard Error, z = z value. Significant p-values are indicated in bold with the following symbols: “*” for p < 0.1, “**” for p < 0.01, and “***” for p < 0.001. See Table S6 for model selection. “Mammal (with domestic)” indicates the results when including domestic mammals (rabbit, sheep, horse, donkey, cat, dog, goat and pig).

**Table S2.** Outputs of the GLMM and GAMM investigating the effect of ambient storage time, read counts, and human pressure defined as the land cover - pollution index (i.e. the site coordinates on the first two axes PC1 and PC2 of the PCA combining CORINE Landcover and heavy metal concentration variables) on alpha diversity (richness). E = Estimate, SE = Standard Error, z = z value. Significant p-values are indicated in bold with the following symbols: “*” for p < 0.1, “**” for p < 0.01, and “***” for p < 0.001. See Table S6 for model selection. “Mammal (with domestic)” indicates the results when including domestic mammals (rabbit, sheep, horse, donkey, cat, dog, goat and pig).

**Table S3.** Outputs of the GLMM and GAMM investigating the effect of ambient storage time, read counts and human pressure defined as the human footprint on alpha diversity (richness). E = Estimate, SE = Standard Error, z = z value. Significant p-values are indicated in bold with the following symbols: “*” for p < 0.1, “**” for p < 0.01, and “***” for p < 0.001. See Table S6 for model selection. “Mammal (with domestic)” indicates the results when including domestic mammals (rabbit, sheep, horse, donkey, cat, dog, goat and pig).

**Table S4.** List of taxa significantly associated to each of the environment types (urban vs rural) using the multilevel pattern analysis on occurrence matrices with the association function (r.g), and a significance level = 0.05 (number of permutations: 999). Significant p-values are indicated in bold with the following symbol: “*” for p < 0.1, “**” for p < 0.01, and “***” for p < 0.001. “Mammal (with domestic)” indicates the results when including domestic mammals (rabbit, sheep, horse, donkey, cat, dog, goat and pig).

**Table S5.** Permutational multivariate analysis of variance (perMANOVA; 999 permutations) investigating the effect of human pressure on community composition for each taxonomic group. Human pressure was defined as A) the urban/rural environment type index. B) the land cover - pollution index. defined as the site coordinates on the first two axes PC1 and PC2 of the PCA combining CORINE land cover and heavy metal concentrations variables. and C) the human footprint index. Significant p-values are indicated in bold with the following symbol: “*” for p < 0.1, “**” for p < 0.01, and “***” for p < 0.001. “Mammal (with domestic)” indicates the results when including domestic mammals (rabbit, sheep, horse, donkey, cat, dog, goat and pig).

**Table S6.** Model selection. AIC and R² values from the GAMM and GLMM richness models of negative binomial family for each taxonomic group (birds, mammals, insects, plants, and fungi), with human pressure defined as A. the urban-rural index (“Environment_type”), B. the land cover-pollution index (“PC1” and “PC2”), and C. the human footprint index (“human_footprint”). Marginal R² values (variance explained by fixed effects) for the GLMMs were calculated using the r2_nakagawa function from the performance package. For GAMMs, the adjusted R² values were extracted from the summary() output (“R-sq.(adj)”). Note that random effects were not included in the models if their variance component was equal to zero according to the GLMM output using the sjPlot::tab_model function (τ00 = 0). Model selection between GLMM and GAMM was based on the lowest AIC, or the highest R² when ΔAIC < 2. The final model is indicated in bold, with its formula detailed in the “FINAL MODEL” column. “Mammal (with domestic)” indicates the results when including domestic mammals (rabbit, sheep, horse, donkey, cat, dog, goat and pig).

**Table S7.** Correlation between the community composition and geographic distance. Mantel tests based on Pearson’s product-moment correlation (mantel function of the vegan package, 999 permutations) investigating the correlation between the community composition (Jaccard dissimilarity distance on presence/absence matrix) and geographic distance (geodesic distance). Significant p-values are indicated in bold with the following symbol: “*” for p < 0.1, “**” for p < 0.01, and “***” for p < 0.001. “Mammal (with domestic)” indicates the results when including domestic mammals (rabbit, sheep, horse, donkey, cat, dog, goat and pig).

## REFERENCES

1. G. Ceballos, P. R. Ehrlich, A. D. Barnosky, A. García, R. M. Pringle, T. M. Palmer, Accelerated modern human–induced species losses: Entering the sixth mass extinction. Science Advances 1, e1400253 (2015).

2. R. H. Cowie, P. Bouchet, B. Fontaine, The Sixth Mass Extinction: fact, fiction or speculation? Biological Reviews 97, 640–663 (2022).

3. J. A. Foley, R. DeFries, G. P. Asner, C. Barford, G. Bonan, S. R. Carpenter, F. S. Chapin, M. T. Coe, G. C. Daily, H. K. Gibbs, J. H. Helkowski, T. Holloway, E. A. Howard, C. J. Kucharik, C. Monfreda, J. A. Patz, I. C. Prentice, N. Ramankutty, P. K. Snyder, Global Consequences of Land Use. Science 309, 570–574 (2005).

4. H. M. Pereira, I. S. Martins, I. M. D. Rosa, H. Kim, P. Leadley, A. Popp, D. P. van Vuuren, G. Hurtt, L. Quoss, A. Arneth, D. Baisero, M. Bakkenes, R. Chaplin-Kramer, L. Chini, M. Di Marco, S. Ferrier, S. Fujimori, C. A. Guerra, M. Harfoot, T. D. Harwood, T. Hasegawa, V. Haverd, P. Havlík, S. Hellweg, J. P. Hilbers, S. L. L. Hill, A. Hirata, A. J. Hoskins, F. Humpenöder, J. H. Janse, W. Jetz, J. A. Johnson, A. Krause, D. Leclère, T. Matsui, J. R. Meijer, C. Merow, M. Obersteiner, H. Ohashi, A. De Palma, B. Poulter, A. Purvis, B. Quesada, C. Rondinini, A. M. Schipper, J. Settele, R. Sharp, E. Stehfest, B. B. N. Strassburg, K. Takahashi, L. Talluto, W. Thuiller, N. Titeux, P. Visconti, C. Ware, F. Wolf, R. Alkemade, Global trends and scenarios for terrestrial biodiversity and ecosystem services from 1900 to 2050. Science 384, 458–465 (2024).

5. Y. Zhou, A. C. G. Varquez, M. Kanda, High-resolution global urban growth projection based on multiple applications of the SLEUTH urban growth model. Sci Data 6, 34 (2019).

6. S. Lokatis, J. M. Jeschke, M. Bernard-Verdier, S. Buchholz, H.-P. Grossart, F. Havemann, F. Hölker, Y. Itescu, I. Kowarik, S. Kramer-Schadt, D. Mietchen, C. L. Musseau, A. Planillo, C. Schittko, T. M. Straka, T. Heger, Hypotheses in urban ecology: building a common knowledge base. Biological Reviews 98, 1530–1547 (2023).

7. T. Christmann, I. Kowarik, M. Bernard-Verdier, S. Buchholz, A. Hiller, B. Seitz, M. von der Lippe, Phenology of grassland plants responds to urbanization. Urban Ecosyst 26, 261– 275 (2023).

8. S. Karan, S. Saraswat, B. S. Anusha, Light pollution and the impacts on biodiversity: the dark side of light. Biodiversity 24, 194–199 (2023).

9. V. Zaffaroni-Caorsi, C. Both, R. Márquez, D. Llusia, P. Narins, M. Debon, M. Borges-Martins, Effects of anthropogenic noise on anuran amphibians. Bioacoustics 32, 90–120 (2023).

10. M. L. McKinney, Effects of urbanization on species richness: A review of plants and animals. Urban Ecosyst 11, 161–176 (2008).

11. A. Perez, S. E. Diamond, Idiosyncrasies in cities: evaluating patterns and drivers of ant biodiversity along urbanization gradients. Journal of Urban Ecology 5, juz017 (2019).

12. E. Carlon, D. M. Dominoni, The role of urbanization in facilitating the introduction and establishment of non-native animal species: a comprehensive review. Journal of Urban Ecology 10, juae015 (2024).

13. S. H. Faeth, C. Bang, S. Saari, Urban biodiversity: patterns and mechanisms. Annals of the New York Academy of Sciences 1223, 69–81 (2011).

14. A. Méndez, T. Montalvo, R. Aymí, M. Carmona, J. Figuerola, J. Navarro, Adapting to urban ecosystems: unravelling the foraging ecology of an opportunistic predator living in cities. Urban Ecosyst 23, 1117–1126 (2020).

15. R. B. Blair, “Birds and Butterflies Along Urban Gradients in Two Ecoregions of the United States: Is Urbanization Creating a Homogeneous Fauna?” in Biotic Homogenization, J. L. Lockwood, M. L. McKinney, Eds. (Springer US, Boston, MA, 2001; 10.1007/978-1-4615-1261-5_3), pp. 33–56.

16. S. Lokatis, J. M. Jeschke, Urban biotic homogenization: Approaches and knowledge gaps. Ecological Applications 32, e2703 (2022).

17. M. L. McKinney, Urbanization as a major cause of biotic homogenization. Biological Conservation 127, 247–260 (2006).

18. M. L. McKinney, Urbanization, Biodiversity, and Conservation: The impacts of urbanization on native species are poorly studied, but educating a highly urbanized human population about these impacts can greatly improve species conservation in all ecosystems. BioScience 52, 883–890 (2002).

19. J. H. Connell, Diversity in Tropical Rain Forests and Coral Reefs. Science 199, 1302–1310 (1978).

20. J. P. Grime, Competitive Exclusion in Herbaceous Vegetation. Nature 242, 344–347 (1973).

21. S. H. Roxburgh, K. Shea, J. B. Wilson, The Intermediate Disturbance Hypothesis: Patch Dynamics and Mechanisms of Species Coexistence. Ecology 85, 359–371 (2004).

22. P. Werner, R. Zahner, “Urban Patterns and Biological Diversity: A Review” in Urban Biodiversity and Design (John Wiley & Sons, Ltd, 2010; https://onlinelibrary.wiley.com/doi/abs/10.1002/9781444318654.ch7), pp. 145–173.

23. L. Celesti-Grapow, P. Pyšek, V. Jarošík, C. Blasi, Determinants of native and alien species richness in the urban flora of Rome. Diversity and Distributions 12, 490–501 (2006).

24. R. Haines-Young, Land use and biodiversity relationships. Land Use Policy 26, S178– S186 (2009).

25. R. Rosa García, M. D. Fraser, R. Celaya, L. M. M. Ferreira, U. García, K. Osoro, Grazing land management and biodiversity in the Atlantic European heathlands: a review. Agroforest Syst 87, 19–43 (2013).

26. N. Müller, M. Ignatieva, C. H. Nilon, P. Werner, W. C. Zipperer, “Patterns and Trends in Urban Biodiversity and Landscape Design” in In: Elmqvist, T., et al. Urbanization, Biodiversity and Ecosystem Services: Challenges and Opportunities: A Global Assessment (Dordrecht, Springer., 2013; 10.1007/978-94-007-7088-1_3), pp. 123–174.

27. P. Reidsma, T. Tekelenburg, M. van den Berg, R. Alkemade, Impacts of land-use change on biodiversity: An assessment of agricultural biodiversity in the European Union. Agriculture, Ecosystems & Environment 114, 86–102 (2006).

28. D. Senapathi, L. G. Carvalheiro, J. C. Biesmeijer, C.-A. Dodson, R. L. Evans, M. McKerchar, R. D. Morton, E. D. Moss, S. P. M. Roberts, W. E. Kunin, S. G. Potts, The impact of over 80 years of land cover changes on bee and wasp pollinator communities in England. Proceedings of the Royal Society B: Biological Sciences 282, 20150294 (2015).

29. C. J. Marsh, E. C. Turner, B. W. Blonder, B. Bongalov, S. Both, R. S. Cruz, D. M. O. Elias, D. Hemprich-Bennett, P. Jotan, V. Kemp, U. H. Kritzler, S. Milne, D. T. Milodowski, S. L. Mitchell, M. M. Pillco, M. H. Nunes, T. Riutta, S. J. B. Robinson, E. M. Slade, H. Bernard, D. F. R. P. Burslem, A. Y. C. Chung, E. L. Clare, D. A. Coomes, Z. G. Davies, D. P. Edwards, D. Johnson, P. Kratina, Y. Malhi, N. Majalap, R. Nilus, N. J. Ostle, S. J. Rossiter, M. J. Struebig, J. A. Tobias, M. Williams, R. M. Ewers, O. T. Lewis, G. Reynolds, Y. A. Teh, A. Hector, Tropical forest clearance impacts biodiversity and function, whereas logging changes structure. Science 387, 171–175 (2025).

30. J. E. Littlefair, J. J. Allerton, A. S. Brown, D. M. Butterfield, C. Robins, C. K. Economou, N. R. Garrett, E. L. Clare, Air-quality networks collect environmental DNA with the potential to measure biodiversity at continental scales. Current Biology 33, R426–R428 (2023).

31. O. Tournayre, J. E. Littlefair, N. R. Garrett, J. J. Allerton, A. S. Brown, M. E. Cristescu, E. L. Clare, First national survey of terrestrial biodiversity using airborne eDNA. bioRxiv [Preprint] (2025). 10.1101/2025.04.07.647580.

32. K. Deiner, H. M. Bik, E. Mächler, M. Seymour, A. Lacoursière-Roussel, F. Altermatt, S. Creer, I. Bista, D. M. Lodge, N. de Vere, M. E. Pfrender, L. Bernatchez, Environmental DNA metabarcoding: Transforming how we survey animal and plant communities. Molecular Ecology 26, 5872–5895 (2017).

33. B. A. Williams, O. Venter, J. R. Allan, S. C. Atkinson, J. A. Rehbein, M. Ward, M. D. Marco, H. S. Grantham, J. Ervin, S. J. Goetz, A. J. Hansen, P. Jantz, R. Pillay, S. Rodríguez-Buriticá, C. Supples, A. L. S. Virnig, J. E. M. Watson, Change in Terrestrial Human Footprint Drives Continued Loss of Intact Ecosystems. One Earth 3, 371–382 (2020).

34. V. Cazalis, Species richness response to human pressure hides important assemblage transformations. Proceedings of the National Academy of Sciences 119, e2107361119 (2022).

35. S. J. Mayor, J. F. Cahill, F. He, P. Sólymos, S. Boutin, Regional boreal biodiversity peaks at intermediate human disturbance. Nat Commun 3, 1142 (2012).

36. I. Kowarik, Novel urban ecosystems, biodiversity, and conservation. Environmental Pollution 159, 1974–1983 (2011).

37. A. P. Hendry, K. M. Gotanda, E. I. Svensson, Human influences on evolution, and the ecological and societal consequences. Philosophical Transactions of the Royal Society B: Biological Sciences 372, 20160028 (2017).

38. J. E. Kemp, A. G. Ellis, Significant Local-Scale Plant-Insect Species Richness Relationship Independent of Abiotic Effects in the Temperate Cape Floristic Region Biodiversity Hotspot. PLoS One 12, e0168033 (2017).

39. M. C. Palmersheim, R. Schürch, M. E. O’Rourke, J. Slezak, M. J. Couvillon, If You Grow It, They Will Come: Ornamental Plants Impact the Abundance and Diversity of Pollinators and Other Flower-Visiting Insects in Gardens. Horticulturae 8, 1068 (2022).

40. K. C. R. Baldock, M. A. Goddard, D. M. Hicks, W. E. Kunin, N. Mitschunas, L. M. Osgathorpe, S. G. Potts, K. M. Robertson, A. V. Scott, G. N. Stone, I. P. Vaughan, J. Memmott, Where is the UK’s pollinator biodiversity? The importance of urban areas for flower-visiting insects. Proceedings of the Royal Society B: Biological Sciences 282, 20142849 (2015).

41. E. Sebastián-González, J. M. Barbosa, J. M. Pérez-García, Z. Morales-Reyes, F. Botella, P. P. Olea, P. Mateo-Tomás, M. Moleón, F. Hiraldo, E. Arrondo, J. A. Donázar, A. Cortés-Avizanda, N. Selva, S. A. Lambertucci, A. Bhattacharjee, A. Brewer, J. D. Anadón, E. Abernethy, O. E. Rhodes Jr, K. Turner, J. C. Beasley, T. L. DeVault, A. Ordiz, C. Wikenros, B. Zimmermann, P. Wabakken, C. C. Wilmers, J. A. Smith, C. J. Kendall, D. Ogada, E. R. Buechley, E. Frehner, M. L. Allen, H. U. Wittmer, J. R. A. Butler, J. T. du Toit, J. Read, D. Wilson, K. Jerina, M. Krofel, R. Kostecke, R. Inger, A. Samson, L. Naves-Alegre, J. A. Sánchez-Zapata, Scavenging in the Anthropocene: Human impact drives vertebrate scavenger species richness at a global scale. Global Change Biology 25, 3005–3017 (2019).

42. R. B. Blair, Land Use and Avian Species Diversity Along an Urban Gradient. Ecological Applications 6, 506–519 (1996).

43. J. M. Parody, F. J. Cuthbert, E. H. Decker, The effect of 50 years of landscape change on species richness and community composition. Global Ecology and Biogeography 10, 305– 313 (2001).

44. A. Trouwborst, P. C. McCormack, E. Martínez Camacho, Domestic cats and their impacts on biodiversity: A blind spot in the application of nature conservation law. People and Nature 2, 235–250 (2020).

45. H. Steinfeld, H. A. Mooney, F. Schneider, L. E. Neville, Livestock in a Changing Landscape, Volume 1: Drivers, Consequences, and Responses (Island Press, 2013).

46. A. Baselga, Partitioning the turnover and nestedness components of beta diversity. Global Ecology and Biogeography 19, 134–143 (2010).

47. A. Baselga, F. Leprieur, Comparing methods to separate components of beta diversity. Methods in Ecology and Evolution 6, 1069–1079 (2015).

48. J. Soininen, J. Heino, J. Wang, A meta-analysis of nestedness and turnover components of beta diversity across organisms and ecosystems. Global Ecology and Biogeography 27, 96–109 (2018).

49. J. P. Wayman, J. P. Sadler, T. A. M. Pugh, T. E. Martin, J. A. Tobias, T. J. Matthews, Identifying the Drivers of Spatial Taxonomic and Functional Beta-Diversity of British Breeding Birds. Front. Ecol. Evol. 9 (2021).

50. J. Clavel, R. Julliard, V. Devictor, Worldwide decline of specialist species: toward a global functional homogenization? Frontiers in Ecology and the Environment 9, 222–228 (2011).

51. M. Vellend, The Biodiversity Conservation Paradox. American Scientist 105, 94–101 (2017).

52. H. Hillebrand, B. Blasius, E. T. Borer, J. M. Chase, J. A. Downing, B. K. Eriksson, C. T. Filstrup, W. S. Harpole, D. Hodapp, S. Larsen, A. M. Lewandowska, E. W. Seabloom, D. B. Van de Waal, A. B. Ryabov, Biodiversity change is uncoupled from species richness trends: Consequences for conservation and monitoring. Journal of Applied Ecology 55, 169–184 (2018).

53. B. M. Fernandez-Going, S. P. Harrison, B. L. Anacker, H. D. Safford, Climate interacts with soil to produce beta diversity in Californian plant communities. Ecology 94, 2007–2018 (2013).

54. F. Zellweger, T. Roth, H. Bugmann, K. Bollmann, Beta diversity of plants, birds and butterflies is closely associated with climate and habitat structure. Global Ecology and Biogeography 26, 898–906 (2017).

55. T.-H. Macher, R. Schütz, T. Hörren, A. J. Beermann, F. Leese, It’s raining species: Rainwash eDNA metabarcoding as a minimally invasive method to assess tree canopy invertebrate diversity. Environmental DNA 5, 3–11 (2023).

56. N. Abrego, V. Norros, P. Halme, P. Somervuo, H. Ali-Kovero, O. Ovaskainen, Give me a sample of air and I will tell which species are found from your region: Molecular identification of fungi from airborne spore samples. Molecular Ecology Resources 18, 511– 524 (2018).

57. F. Roger, H. R. Ghanavi, N. Danielsson, N. Wahlberg, J. Löndahl, L. B. Pettersson, G. K. S. Andersson, N. Boke Olén, Y. Clough, Airborne environmental DNA metabarcoding for the monitoring of terrestrial insects—A proof of concept from the field. Environmental DNA 4, 790–807 (2022).

58. R. M. Harrison, D. Laxen, S. Moorcroft, K. Laxen, Processes affecting concentrations of fine particulate matter (PM2.5) in the UK atmosphere. Atmospheric Environment 46, 115– 124 (2012).

59. P. D. Lamb, E. Hunter, J. K. Pinnegar, S. Creer, R. G. Davies, M. I. Taylor, How quantitative is metabarcoding: A meta-analytical approach. Molecular Ecology 28, 420–430 (2019).

60. M. A. Lopes, S. F. Ferrari, Effects of Human Colonization on the Abundance and Diversity of Mammals in Eastern Brazilian Amazonia. Conservation Biology 14, 1658–1665 (2000).

61. S. Saari, S. Richter, M. Higgins, M. Oberhofer, A. Jennings, S. H. Faeth, Urbanization is not associated with increased abundance or decreased richness of terrestrial animals - dissecting the literature through meta-analysis. Urban Ecosyst 19, 1251–1264 (2016).

62. M. C. Yates, T. M. Wilcox, S. Kay, P. Peres-Neto, D. D. Heath, A Framework to Unify the Relationship Between Numerical Abundance, Biomass, and Environmental DNA. Environmental DNA 7, e70073 (2025).

63. A. O. Shelton, Z. J. Gold, A. J. Jensen, E. D′Agnese, E. Andruszkiewicz Allan, A. Van Cise, R. Gallego, A. Ramón-Laca, M. Garber-Yonts, K. Parsons, R. P. Kelly, Toward quantitative metabarcoding. Ecology 104, e3906 (2023).

64. J. Battin, When Good Animals Love Bad Habitats: Ecological Traps and the Conservation of Animal Populations. Conservation Biology 18, 1482–1491 (2004).

65. J. Zuñiga-Palacios, I. Zuria, I. Castellanos, C. Lara, G. Sánchez-Rojas, What do we know (and need to know) about the role of urban habitats as ecological traps? Systematic review and meta-analysis. Science of The Total Environment 780, 146559 (2021).

66. K. R. Williams, E. C. Braysher, J. H. L. Cheong, C. C. Robins, V. Kantilal, D. M. Butterfield, A. Lilley, C. Bradshaw, A. S. Brown, R. J. C. Brown, “Annual Report for 2023 on the UK Heavy Metals Monitoring Network” (55, National Physical Laboratory, Atmospheric Environmental Science Department, 2024); 10.47120/npl.ENV55.

67. J. Oksanen, G. L. Simpson, F. G. Blanchet, R. Kindt, P. Legendre, P. R. Minchin, R. B. O’Hara, P. Solymos, M. H. H. Stevens, E. Szoecs, H. Wagner, M. Barbour, M. Bedward, B. Bolker, D. Borcard, G. Carvalho, M. Chirico, M. D. Caceres, S. Durand, H. B. A. Evangelista, R. FitzJohn, M. Friendly, B. Furneaux, G. Hannigan, M. O. Hill, L. Lahti, D. McGlinn, M.-H. Ouellette, E. R. Cunha, T. Smith, A. Stier, C. J. F. T. Braak, J. Weedon, vegan: Community Ecology Package, version 2.6-8 (2024); https://cran.r-project.org/web/packages/vegan/index.html.

68. B. Cole, B. De la Barreda, A. Hamer, T. Codd, M. Payne, L. Chan, G. Smith, H. Balzter, Corine land cover 2018 for the UK, Isle of Man, Jersey and Guernsey., NERC EDS Environmental Information Data Centre. (2021); 10.5285/084e0bc6-e67f-4dad-9de6-0c698f60e34d.

69. © Crown, © Crown copyright Defra via uk-air.defra.gov.uk, licenced under the Open Government Licence (OGL)., (2025).

70. R Core Team, R: A language and environment for statistical computing, R Foundation for Statistical Computing (2024); https://www.R-project.org/.

71. M. E. Brooks, K. Kristensen, K. J. van Benthem, A. Magnusson, C. W. Berg, A. Nielsen, H. J. Skaug, M. Maechler, B. M. Bolker, glmmTMB Balances Speed and Flexibility Among Packages for Zero-inflated Generalized Linear Mixed Modeling. The R Journal 9, 378–400 (2017).

72. S. N. Wood, Fast stable restricted maximum likelihood and marginal likelihood estimation of semiparametric generalized linear models. Journal of the Royal Statistical Society (B) 73, 3–36 (2011).

73. G. L. Simpson, gratia: An R package for exploring generalized additive models. Journal of Open Source Software 9, 6962 (2024).

74. D. Lüdecke, ggeffects: Tidy Data Frames of Marginal Effects from Regression Models. Journal of Open Source Software 3, 772 (2018).

75. M. De Cáceres, P. Legendre, Associations between species and groups of sites: indices and statistical inference. Ecology 90, 3566–3574 (2009).

76. S. Dray, D. Bauman, G. Blanchet, D. Borcard, S. Clappe, G. Guenard, T. Jombart, G. Larocque, P. Legendre, N. Madi, H. H. Wagner, A. Siberchicot, adespatial: Multivariate Multiscale Spatial Analysis, version 0.3-24 (2024); https://cran.r-project.org/web/packages/adespatial/index.html.

77. J. B. Kruskal, Multidimensional scaling by optimizing goodness of fit to a nonmetric hypothesis. Psychometrika 29, 1–27 (1964).

